# Mutational scanning reveals oncogenic *CTNNB1* mutations have diverse effects on signalling and clinical traits

**DOI:** 10.1101/2023.11.09.566307

**Authors:** Anagha Krishna, Alison Meynert, Martijn Kelder, Ailith Ewing, Shahida Sheraz, Anna Ferrer-Vaquer, Graeme Grimes, Hannes Becher, Ryan Silk, Colin A Semple, Timothy Kendall, Anna-Katerina Hadjantonakis, Tom Bird, Joseph A Marsh, Peter Hohenstein, Andrew J Wood, Derya D Ozdemir

## Abstract

*CTNNB1*, the gene encoding β-catenin, is a frequent target for oncogenic mutations activating the canonical Wnt signalling pathway, typically via missense mutations within a degron hotspot motif in exon 3. Here, we combine saturation genome editing with a fluorescent reporter assay to quantify signalling phenotypes for all 342 missense mutations in the mutation hotspot, including 74 recurrent mutations reported in over 6000 tumours. Our data define the genetic requirements for β-catenin degron function and reveal diverse levels of signal activation among known driver mutations. Tumorigenesis in different human tissues involves selection for *CTNNB1* mutations spanning distinct ranges of effect size. In hepatocellular carcinoma, mutations that activate β-catenin relatively weakly are associated with worse prognosis compared to stronger activating mutations, despite greater immune cell infiltration in the tumour microenvironment. Our work therefore provides a resource to understand mutational diversity within a pan-cancer mutation hotspot, with potential implications for targeted therapy.

## Main Text

The canonical Wnt signalling pathway is essential for normal development and is erroneously activated in many types of cancer (Clevers and Nusse 2012). The main intracellular signal transducer is β-catenin, encoded by the *CTNNB1* gene. Under normal conditions the level of β-catenin available for signalling is kept low by a destruction complex consisting of APC, AXIN2, CK1 and GSK3β. In the presence of Wnt ligands or mutations that reduce destruction complex function, β-catenin accumulates, enters the nucleus and activates target genes via association with TCF transcription factors (Figure 1A).

**Figure 1:**
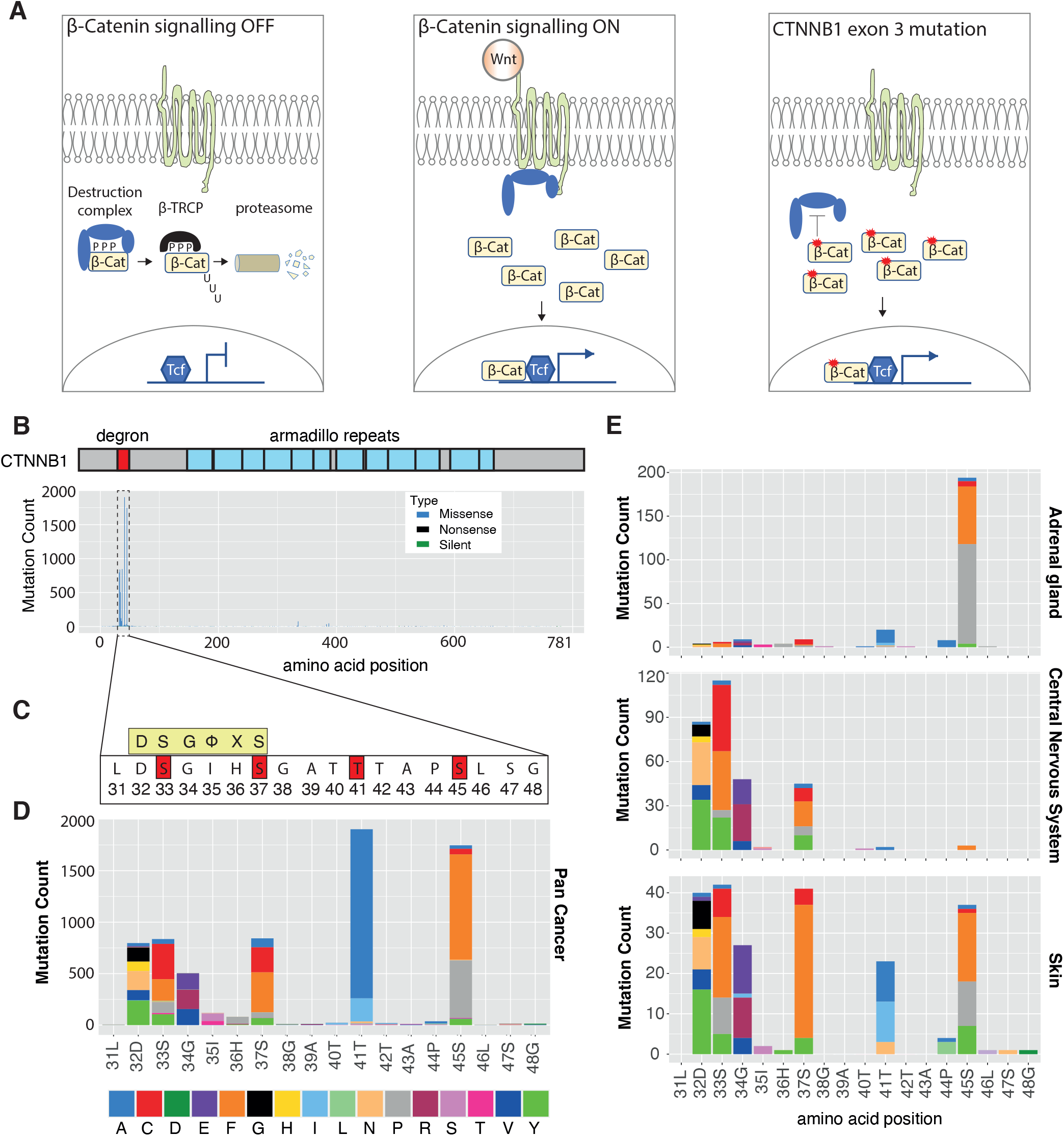
Tissue-specific mutation profiles at the *CTNNB1* degron in human cancer. **A.** Simplified schematic of Wnt pathway regulation in normal development and in the presence of exon 3 degron mutations. (Left) In the absence of Wnt ligand, the destruction complex phosphorylates sites within the exon 3 degron to allow docking of the E3 ligase receptor β-TRCP, followed by ubiquitination and degradation via the proteasome. (Centre) In the presence of Wnt ligand, the destruction complex is sequestered at the cell membrane. ß-Catenin accumulates and translocates to the nucleus where it associates with Tcf transcription factors to activate target genes. (Right) Exon 3 mutations prevent destruction complex activity to uncouple ß-Catenin accumulation from Wnt ligand binding, locking the pathway in the active state. **B.** Histogram showing the frequency of mutations at each amino acid position of human *CTNNB1* across all tumours present in the COSMIC database. Blue boxes represent the positions of armadillo repeat domains, and the dashed rectangle indicates the region blown up in panels C & D. **C**. The amino acid sequence of the *CTNNB1* mutation hotspot. The consensus docking site for the β-TRCP E3 ligase substrate receptor is shown in a yellow box, where ϕ indicates any hydrophobic amino acid and X indicates any amino acid. Phosphorylation sites known to be critical for CTNNB1 turnover are highlighted in red boxes. **D**. Histogram shows the frequency of each amino acid substitution across each position in the *CTNNB1* mutation hotspot, using data from all tumours in the COSMIC database. **E.** Histograms show distinct distributions of mutations within the *CTNNB1* mutation hotspot for COSMIC tumours of the adrenal gland, central nervous system and skin. Data from other primary tissue sites are shown in Figure S1.

*APC* loss-of-function mutations occur in most colorectal cancers. Gain-of-function changes within a degron motif in *CTNNB1* exon 3 are among the most common mutations in cancers of the liver and endometrium and occur frequently in tumours originating in several other tissues (Çelen et al. 2015). A large body of work has demonstrated that mutations within this pan-cancer hotspot region act in a dominant manner by blocking destruction complex activity (reviewed in (Gao et al. 2018)). Positions 32 – 37 of CTNNB1 comprise the DpSGXXpS docking motif for the E3 ubiquitin ligase substrate receptor β-TRCP, where pS denotes phosphoserine and X any residue (Wu et al. 2003). The phospho-regulatory cascade involves an initial CK1-mediated phosphorylation at S45 (‘CK1 priming’), followed by GSK3β -mediated phosphorylation at positions T41, S37 and S33 to enable β-TRCP docking.

Analysis of *APC* mutations in patient tumour material and mouse models has shown that tumours often select for mutations that result in a specific ‘just-right’ level of β-catenin signalling (Albuquerque et al. 2002; Pollard et al. 2009; Feng et al. 2015). However, although analysis of a limited number of mutations within *CTNNB1* has suggested that different mutations can activate to different degrees (Rebouissou et al. 2016), how the full spectrum of *CTNNB1* mutational diversity impacts on signalling and tumorigenesis is unknown.

### *CTNNB1* hotspot mutational patterns are tissue specific

In order to catalogue the complexity of β-catenin mutations in human cancer, we analysed 9248 tumours with *CTNNB1* mutations in the COSMIC database (Figure 1B, Table S1). As previously reported (Çelen et al. 2015), the majority of mutations (86%) were missense substitutions. Of these, 88% occurred in the L31-S48 hotspot region encoded by exon 3 (Figure 1C & 1D, Table S1). Residues T41 and S45 were mutated most frequently in this pan-cancer analysis, comprising 27% and 25% of all hotspot missense mutations, followed by S37, S33, D32 and G34 (Figure 1D).

We next analysed the distribution of missense mutations in the hotspot region within cancers originating from different primary tissue sites (Figure 1E, Figure S1). Tumours associated with some primary sites tended to favour mutations at particular codon positions within the hotspot. For example, S45 mutations are greatly enriched in tumours originating in the adrenal gland and kidney, whereas mutations within the β-TCRP recognition motif were favoured in the central nervous system (Figure 1E). Other tissues such as the liver and skin showed a comparatively broad profile, with mutations distributed across each of the 6 most frequently mutated positions. At individual positions, we also observed biases in the specific amino acid substitutions that were observed across tumour sites. For example, 99% of soft tissue tumours with T41 mutations (1201/1217) carried T41A whereas in pituitary gland, 89% of T41 mutations (77/87) were T41I (Figure S1). Overall, it is clear that there is substantial diversity in the mutational patterns seen in tumours within the *CTNNB1* hotspot.

### Mutational scanning reveals diverse consequences of *Ctnnb1* hotspot mutations

To systematically measure the consequences of *CTNNB1* hotspot mutations on β-catenin signalling, we devised a multiplexed CRISPR homology-directed repair assay to quantify the phenotypes resulting from all 342 possible single amino acid substitutions across positions 31-48. This assay allows phenotypic analysis of variants expressed from the endogenous mouse *Ctnnb1* locus (Figure S2A & S2B, Methods), which is 100% conserved with human at the amino acid level throughout exon 3. The function of each variant was measured in parallel, under normal regulatory control (Findlay et al. 2014; 2018; Buckley et al. 2023), to provide a sensitive and comprehensive assessment of hotspot mutant phenotypes. A library of homology-directed repair (HDR) templates was synthesised, each encoding a single amino acid substitution within the hotspot and flanking synonymous nucleotide changes to enable selective PCR amplification of edited alleles (Figure S2; (Findlay et al. 2014; 2018)).

To establish a system in which the phenotype of individual hotspot mutations could be measured in single cells, we derived primary embryonic stem cells (ESCs) from the Tcf/Lef::H2B-GFP transgenic mouse line (Figure 2A, Figure S2A) (Ferrer-Vaquer et al. 2010; Czechanski et al. 2014) then replaced exons 3 – 6 of *Ctnnb1* with a negative counter selection cassette on one allele. By selectively targeting and replacing the selection cassette with a multiplexed HDR template library in combination with Fluorescence Activated Cell Sorting (FACS – Figure 2B), this allowed us to efficiently generate and measure the functional consequences of different *Ctnnb1* hotspot mutations in a single experimental workflow (Figure 2C, full procedure described in Figure S2 and Methods).

**Figure 2:**
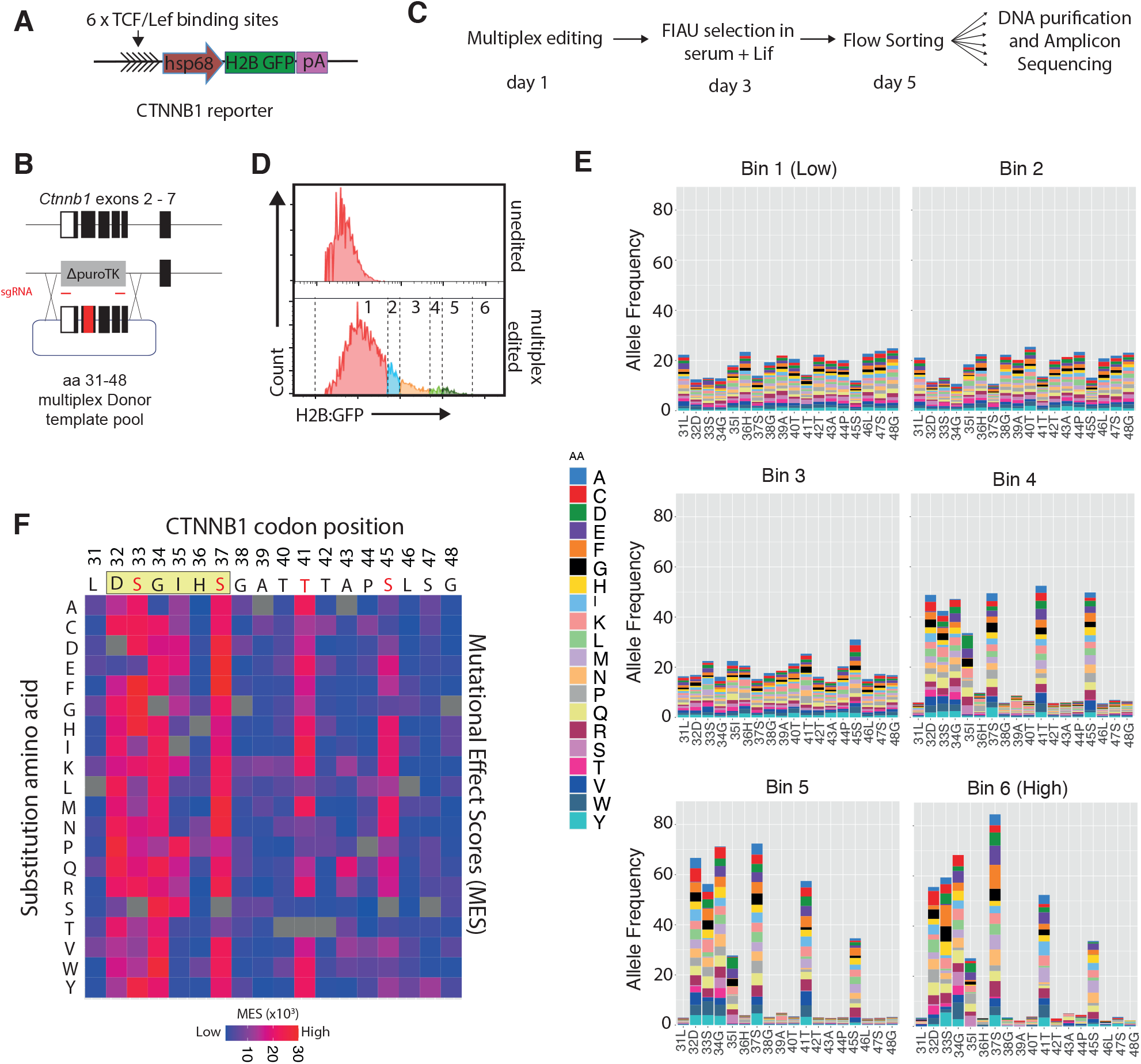
Systematic measurement of β-Catenin activation via mutational scanning. **A.** The CTNNB1/TCF reporter construct described in (Ferrer-Vaquer et al 2010). **B.** Editing strategy used to achieve monoallelic multiplexed homology-directed repair. As shown in Figure S4, primary murine ESCs were engineered such that exons 2 – 6 of *CTNNB1* were heterozygously replaced with a puroTK cassette. Two sgRNAs then cleave at either flank of this integrated cassette while leaving the wildtype allele unmodified. High throughput gene synthesis was used to generate double stranded DNA molecules encoding each individual amino acid substitution across positions 31 – 48 (n = 342 substitutions), which were then cloned into a vector containing 1kb homology arms on either side. **C.** Timeline for the multiplex editing assay to classify *CTNNB1* hotspot mutations. Further details of panels A – C are provided in Figure S4 and in the Materials and Methods section. **D.** Flow cytometry histograms of cells subjected to the steps indicated in panel C, with (bottom) versus without (top) multiplex editing. **E.** The frequency of individual missense mutations across each position in the mutation hotspot in cell populations sorted according to the scheme shown in panel D. The colour scheme used to represent different missense mutations is shown to the right. **F.** Heatmap representation of mutational effect scores (MES), showing the average activity profile for every possible amino acid substitution at the *CTNNB1* mutation hotspot, reconstructed from data shown in panel E. Individual scores are provided in Table S2. The consensus motif for ß-TRCP docking is shown in a beige rectangle above the heatmap, with known phosphorylation sites highlighted in red.

After editing and selection, a subset of cells in the population expressed the Tcf/Lef::H2B-GFP reporter at elevated levels (Figure 2D), consistent with Wnt pathway activation. Cells were sorted into six equally log-spaced bins (referred to as “bin P1-P6”) based on increasing reporter GFP signal (Figure 2D), and genomic DNA was purified and subjected to amplicon deep sequencing across the *Ctnnb1* hotspot region using primers that selectively amplified HDR-edited alleles (Figure S2). In addition, amplicon sequencing was conducted on control samples including the untransfected HDR donor template library (“plasmid”), and on genomic DNA from cells that had been edited but not separated based on GFP expression (“pool”, Figure S3). This allowed us to enumerate the frequency of *Ctnnb1* alleles associated with each range of reporter gene activity relative to their frequency in the total cell population. The resulting codon replacement frequencies showed high correlations between biological replicates (Pearson’s R, range 0.65 – 0.98), and were therefore merged for downstream analysis. Codon replacement frequencies also correlated well between plasmid and pool samples (Pearson’s R=0.63), indicating that HDR efficiency was consistent and unbiased throughout the targeted region (Figure S3).

Whereas the replacement frequency was relatively constant across positions in the unsorted cell population (Figure S3), it varied substantially across cells expressing different levels of GFP (Figure 2E), confirming that different hotspot mutation patterns can lead to different levels of β-catenin activity. The codons with high mutation frequencies in tumours (D32, S33, G34, S37, T41 and S45) were found at low frequency in low activity bins, but made up a large proportion of sequences in bins of higher activity (Figure 2E). Mutations at these sites are known to disrupt β-catenin phospho-regulation and increase Wnt signalling (Morin et al. 1997; Korinek et al. 1997), confirming the utility of this system to functionally classify *CTNNB1* mutations.

We next reconstructed an average activity profile for each amino acid substitution by combining the codon replacement frequency in each of the 6 bins with the average fluorescence value of cells in the corresponding bin (Supplemental methods, (Kosuri et al. 2013)). The resulting mutational effect scores (MES) provided a metric to compare the phenotypic consequences of different amino acid substitutions within the hotspot region to gain insight into β-catenin regulation (Figure 2F, Table S2).

To evaluate the quality of this dataset, we compared the MES values with scores produced by 50 different computational variant effect predictors (VEPs) for the same substitutions (Table S3, (Livesey and Marsh 2023)). The strongest correlation with MES values was observed for EVE, a recently developed method that uses deep generative models to calculate variant effect scores from evolutionary sequence information (Frazer et al. 2021), and was recently shown to be a top-performing VEP when benchmarked against deep mutational scanning data and clinical variants (Livesey and Marsh 2023). Notably, the correlation between the MES values for the *CTNNB1* mutation hotspot and the corresponding EVE predictions (Spearman’s ρ=0.662, Figure S4A, S4B) was higher than observed for any other protein in this benchmarking study, and higher than observed for all but 3 out of 1430 VEP/DMS comparisons (Livesey and Marsh 2023). Thus, the strong correlation we observe here is supportive of the high quality of the current dataset.

Notably, EVE predicted mutations at two codon positions (G38 and P44) to elicit substantially stronger phenotypes relative to the MES values (Figure S4B, S4C). However, neither codon is frequently mutated in human cancer (Figure 1, Figure S1), as would be expected if mutation at these sites activated the canonical WNT pathway. Thus our experimental mutational scanning strategy generates data that are broadly consistent with the best computational predictors, but also appear to be a more reliable guide to the roles of *CTNNB1* mutations in cancer.

### Genotype/phenotype correlations within the *CTNNB1* hotspot

Although mutational scanning scored all 19 possible amino acid substitutions at each position, only a subset can be reached via a single nucleotide change in the human genome (105/342, median = 6 amino acid changes per site, Figure 3A). As expected, this single nucleotide group included almost all (>98%) of the single amino acid substitutions observed in the COSMIC database. Also as expected given the dual specificity of CK1 and GSK3β for phosphorylation at serine and threonine residues, S45T and T41S substitutions yielded low mutational effect scores relative to other substitutions at the same position (Figure 2F, Figure 3B).

**Figure 3:**
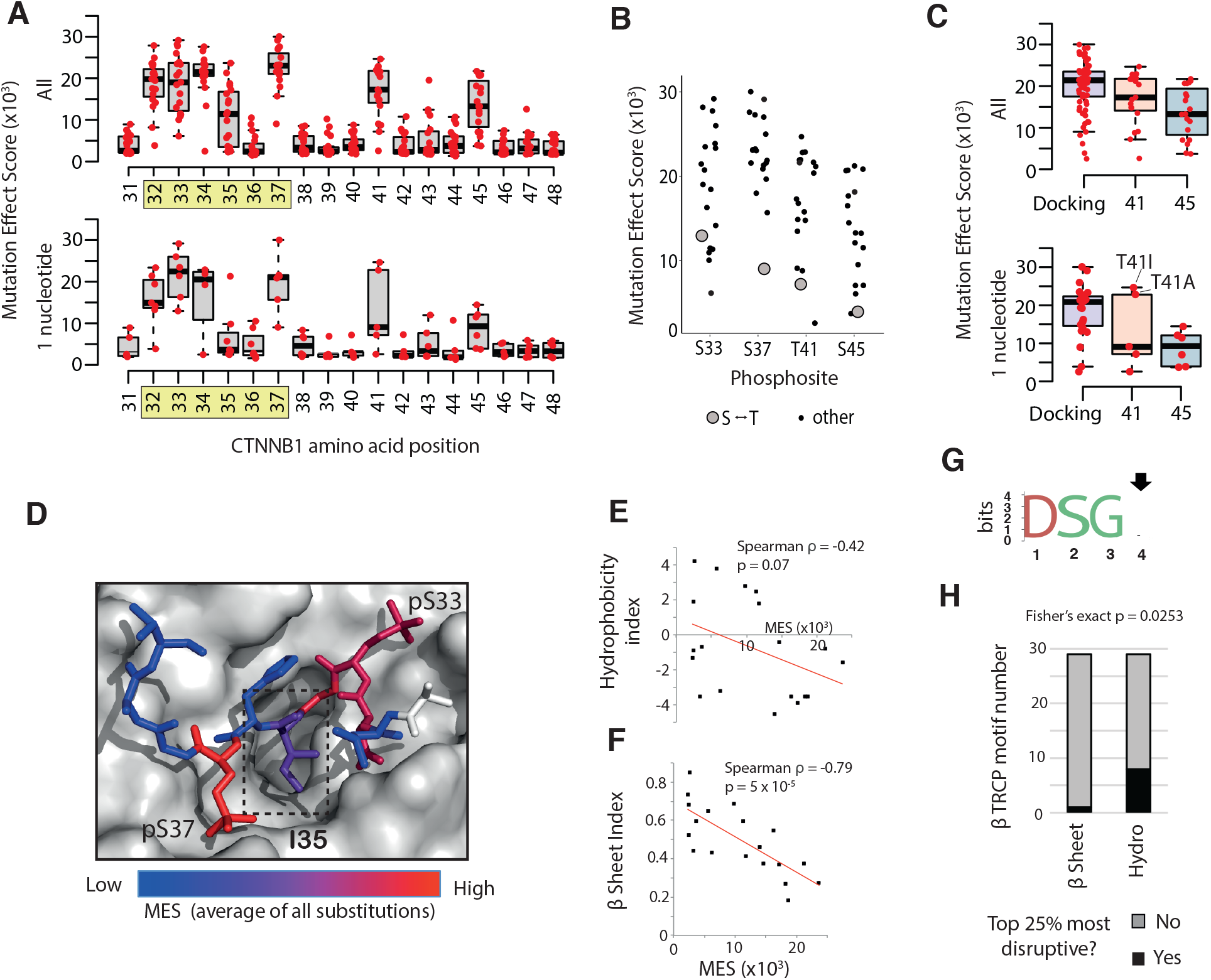
Genotype / phenotype correlations at the *CTNNB1* mutation hotspot. **A.** The distribution of MES values is shown by codon position for all 19 missense substitutions (top), or just the subset that can be reached by a single nucleotide mutation in the human genome (bottom). The core ß-TRCP docking motif is highlighted in yellow. For all boxplots, boxes show the upper and lower quartiles and whiskers show the range. Horizontal lines show the median value, boxes show the upper and lower quartiles and whiskers show the range. **B**. The distribution of MES values for all amino acid substitutions is shown for individual phosphosites, with Serine/Threonine substitutions highlighted. **C.** The distribution of all MES values (top), or just the single nucleotide subset (bottom) for invariable positions within the ß-TRCP docking motif (combined for D32, S33, G34, S37: ‘Docking’), T41, and the CK1 target site at position 45. The two most common T41 mutations in human cancer are labelled in the bottom panel. **D.** Structure of human ß-TRCP (shown in surface mode) in complex with the phosphorylated degron peptide of ß-Catenin (amino acids 30 – 40 are shown). Individual ß-Catenin residues are coloured according to the mean MES value across all substitutions at that position. The box indicates position I35 **E**. Correlation between MES values for individual substitutions at position I35 with the Kyte/Doolittle hydrophobicity scale (Kyte and Doolittle 1982). **F.** Correlation between MES values for individual substitutions at position I35 with the Chou/Fasman ß sheet scale (Chou and Fasman 1978). **G**. Amino Acid logo generated from n = 29 high confidence ß-TRCP docking sites containing the ‘DSGX’ motif (Low et al. 2014). Position 4, equivalent to I35 in ß-Catenin, is highlighted with an arrow. **H**. Stacked bar chart showing the proportion of high confidence ß-TRCP motifs (n = 29) with position 4 residues that rank in the top 25% most disruptive based on the hydrophobicity and ß-sheet indices detailed above.

Prior studies have suggested that S45 mutations activate β-Catenin signalling only weakly, whereas T41 mutations activate moderately and mutations within the core β-TRCP docking motif (32/33/34/37) have strong effects (Rebouissou et al. 2016). In agreement with this we found that mutations at position 45 tended to activate reporter gene expression to a lower level, on average, compared to 32/33/34/37, and this was true both for the single nucleotide mutation group and when all possible mutations were considered (Figure 3C). In particular, substitution of S45 for small amino acids (alanine, glycine, cysteine) was well tolerated, despite removing the CK1 target site thought to be required for N-terminal phosphorylation events and turnover (Amit et al. 2002). It has previously been reported that in-frame deletions at S45 still enable phosphorylation to occur at T41, S37 and S33 in colon cancer cells (Wang et al. 2003; Parker et al. 2020). Collectively, the data therefore suggest that the absence of a large side chain at position S45 enables GSK3β-mediated phosphorylation without CK1 priming.

In our dataset, mutations at T41 had effects that were intermediate, on average, compared with mutations at S45 or within the β-TRCP docking motif, in agreement with previous studies (Rebouissou et al. 2016). However, essentially all cancer mutations at T41 are substitutions for alanine (A) or isoleucine (I). These strong T41 substitutions activate β-Catenin more than most substitutions at positions 32/33/34/37, including several found commonly in cancer (e.g. D32V, D32Y, D32H, S33P, G34R). It is therefore evident that different substitutions at the same position can elicit markedly different effects on β-Catenin signalling. This extends our understanding of *CTNNB1* mutational diversity, with potential implications for patient stratification.

The I35 residue occupies a central position in the docking motif that directly contacts β-TRCP, which has been suggested to require a hydrophobic amino acid side chain (Wu et al. 2003; Çelen et al. 2015). Consistent with this, the I35 side chain can be seen to extend into a cavity on the β-TRCP surface in the co-crystal structure (Figure 3D, (Wu et al. 2003)). MES values for different substitutions at I35 were broadly distributed (Figure 2F, Figure 3B), and showed the expected negative correlation with a hydrophobicity index at this position (Spearman’s ρ=-0.42, Figure 3E, (Kyte and Doolittle 1982)). However, substitution of isoleucine for several polar amino acids (e.g. threonine, asparagine, tyrosine) yielded low MES values (Figure 2F), showing that hydrophobic side chains are not necessary at this position for normal degron function. To investigate this further, we used the AAindex

database (Kawashima et al. 2008) to search for other physicochemical and biochemical amino acid properties that better explained the MES index at I35. Of 566 indices, we observed that those with the strongest correlations were nearly all related to secondary structure propensities. For example, a Spearman correlation of -0.79 was observed with the “normalised frequency of beta sheet” index (Figure 3F, (Chou and Fasman 1978)). While residue I35 does not form a β-sheet specifically in any of the available crystal structures, its dihedral angles place it clearly in the β region of a Ramachandran plot.

To ask whether this secondary structure requirement extended to other β-TRCP docking sites in the proteome, we compiled a list of 28 high confidence β-TRCP-dependent degron sequences containing an exact match to the “DSGX” motif spanning β-Catenin positions 32 – 35 ((Table S4, (Low et al. 2014)). As expected, amino acid identity at position 4 of this motif (corresponding to I35 in β-Catenin) was variable across substrates (Figure 3G). Numerous substrates (8/28) had position 4 amino acids that ranked in the 25% most disruptive on the hydrophobicity scale, whereas only 1/28 featured in the same bracket of the β -sheet scale (Fisher’s exact p = 0.0248). Thus, it appears that requirement for a specific extended backbone conformation can better explain the effects of substitutions at I35 of β-Catenin, and potentially equivalent positions of other β-TRCP docking sites, than the hydrophobicity of the side chain.

### Tissue-specific *CTNNB1* mutation patterns are driven by selection for optimal levels of Wnt signalling

There are notably different frequencies of individual *CTNNB1* hotspot mutations in different tumour types (Figure 1D, Figure S1), and this may reflect a variety of factors. Stem cells in different tissues are subject to different genotoxic insults, which affect overall mutational spectra (Alexandrov et al. 2013; Blokzijl et al. 2016), and could thus affect the probability that particular nucleotide substitutions encoding missense mutations within the *CTNNB1* hotspot become available to selection. Alternatively, or in addition, different tissue environments might favour selection for missense mutations causing levels of activation that are optimal, or ‘just-right’, for their spatio-temporal context (Albuquerque et al. 2002). The current data provide an unusual opportunity to assess the roles of mutational bias and selection in the hotspot variants that accumulate during tumourigenesis.

It has previously been shown that mutational probability, calculated from background rates of nucleotide substitution, is a poor predictor of *CTNNB1* mutation patterns in cancer (Muiños et al. 2021), suggesting a strong influence of selection. To extend these observations, we computed ‘Mutational Likelihood Scores’ (MLS) for each of 342 hotspot missense mutations, using the background rates of nucleotide substitution in Hepatocellular Carcinoma and Endometrial Carcinoma whole exome sequencing (WES) data from TCGA (Figure S5A, S5B, S5C). These MLS scores represent the probability of an amino acid substitution, given the dominant mutational biases seen genome-wide in coding sequences of *CTNNB1*-mutant tumours. MLS values correlated positively across tissues (Pearson R = 0.920 for all mutations & 0.655 for the single nucleotide group, Figure 4A), but were not predictive of the observed mutation frequency in the corresponding tumour type (Figure 4B, negative binomial GLM, no improvement over null model, p_LRT_=0.77 and 0.60, respectively). The data therefore do not support the hypothesis that tissue-specific differences in observed *CTNNB1* mutation patterns are primarily driven by the rates at which the specific mutations arise.

**Figure 4:**
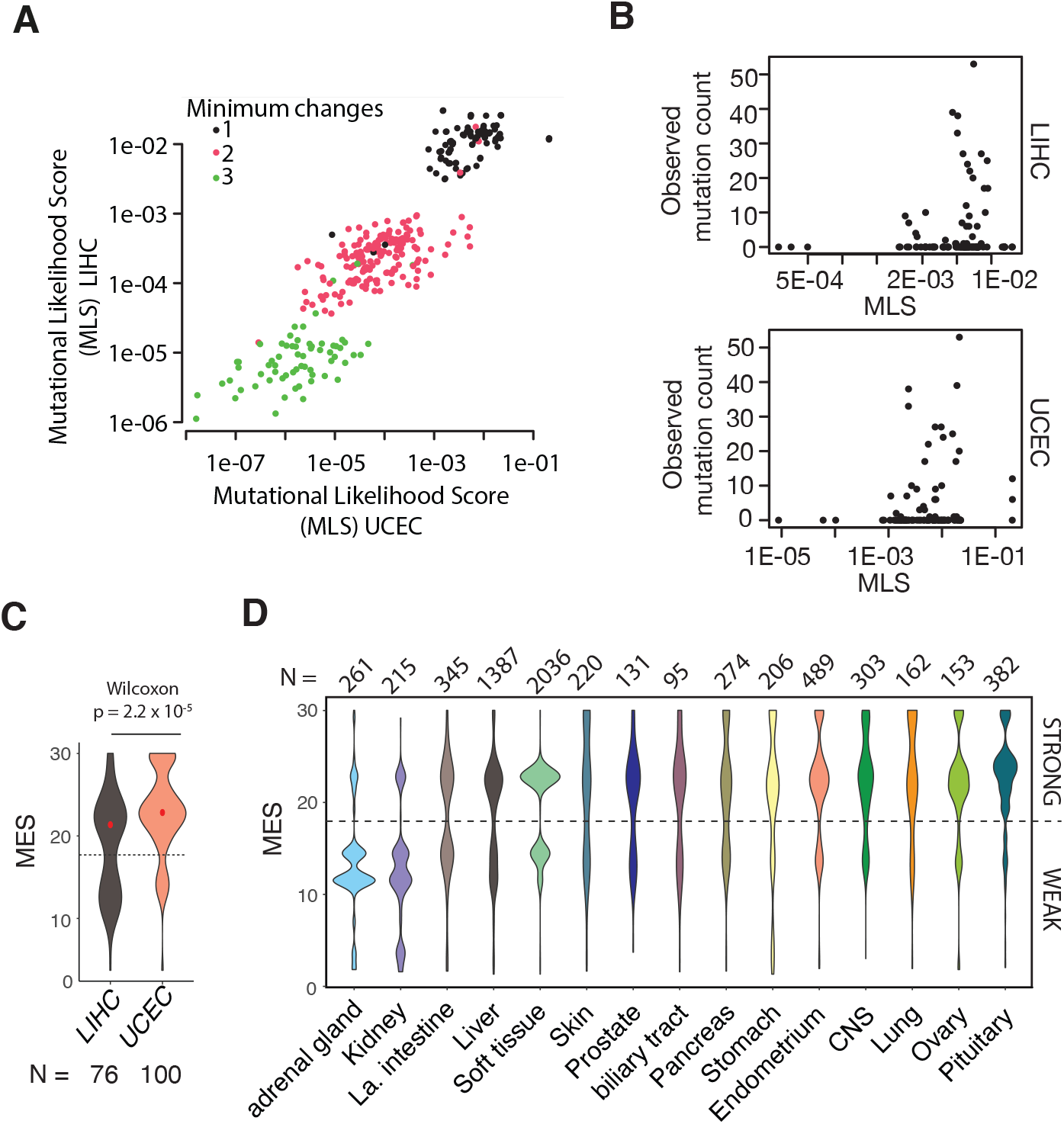
Tissue-specific *CTNNB1* mutation patterns are driven by selection for optimal levels of Wnt signalling: **A.** The relationship between Mutational Likelihood Scores (see Figure S5) in exomes of Hepatocellular Carcinoma (LIHC) and Endometrial Carcinoma (UCEC) tumours from The Cancer Genome Atlas. Scores are shown for all 342 possible amino acid changes, coloured according to the minimum number of nucleotide substitutions required. **B.** The relationship between the observed frequency of specific *CTNNB1* hotspot amino acid substitutions and Mutational Likelihood Scores (MLS) calculated from LIHC and UCEC exomes (See Figure S5). Only amino acid substitutions that can be reached via a single nucleotide mutation (>98% of observed mutations) are shown. **C.** The distribution of MES values in all LIHC and UCEC tumours from TCGA with *CTNNB1* exon 3 hotspot mutations. The dashed line shows the arbitrary threshold used to distinguish STRONG versus WEAK effect mutations in Figure 5. **D.** The distribution of MES values in tumours from different primary sites in the COSMIC dataset. All tissue sites with >100 *CTNNB1* exon 3 mutations are shown. Tissues are ordered according to the ratio of WEAK to STRONG hotspot mutations.

We next asked whether tumours arising in different tissues tend to favour hotspot mutations that activate ß-Catenin signalling to different degrees. This proved to be the case. Mutations in the Endometrial Carcinoma cohort from TCGA (UCEC) were enriched for higher MES values (Mann Whitney p = 2.2 x 10^-5^), whereas in the Hepatocellular Carcinoma cohort (LIHC), mutations spanned a broader range (Figure 4C). Given that mutation bias does not explain the tissue-specific difference in observed mutation frequencies, we postulate they most likely arise via natural selection for different optimal levels of Wnt signalling (Albuquerque et al. 2002).

Even greater variation in MES values was observed in the larger COSMIC dataset (Table S1) across a range of *CTNNB1*-mutant tumours from different primary tissue sites (Figure 4D, One-way ANOVA p < 2 x 10^-19^). Tissues could be divided into categories based on the MES distribution: those which favoured mutations of relatively high effect (e.g. CNS, Pituitary), relatively low effect (e.g. Kidney, Adrenal Gland), or a broad or bimodal distribution (Figure 4C). Bimodal distributions were observed in tissues such as the Large Intestine and Liver, and comprised a lower effect group with mutations at S45 and weaker mutations within the β-TRCP docking motif (e.g. H36P, D32Y, S33P), and a higher effect group containing mutations at S33, G34, S37 And T41. These distributions highlight the potential for wide phenotypic variation, even in tumours with the same site of origin, as a result of mutations within the same *CTNNB1* hotspot region.

### Clinical and molecular correlates of *CTNNB1* mutation strength in Hepatocellular Carcinoma

To gain insight into the effect of *CTNNB1* mutation strength on inter-tumour heterogeneity, we identified Hepatocellular Carcinoma (HCC) samples from TCGA (Cancer Genome Atlas Research Network 2017) with missense mutations in the exon 3 hotspot and available RNA-Seq data, and stratified them into STRONG (n = 47) and WEAK (n = 33) categories based on the bimodal distribution of MES scores observed in this cohort (Figure 4C). Strikingly, survival was significantly poorer for tumours with WEAK mutations compared to those with either STRONG mutations (Figure 5A), or no *CTNNB1* pathway mutation (No mutation, n = 217, Figure 5B). Without stratification by mutation strength, survival of HCC patients with any hotspot missense mutation (n = 80) was not significantly different from the no mutation group (Figure S6A). Moreover, stratification of the exon 3 hotspot mutant group using scores generated by the EVE *in silico* prediction tool ((Frazer et al. 2021), Figure S3), rather than experimental measurements, did not identify significant differences in survival associated with mutations of predicted weak versus strong effect (Figure S6B).

**Figure 5:**
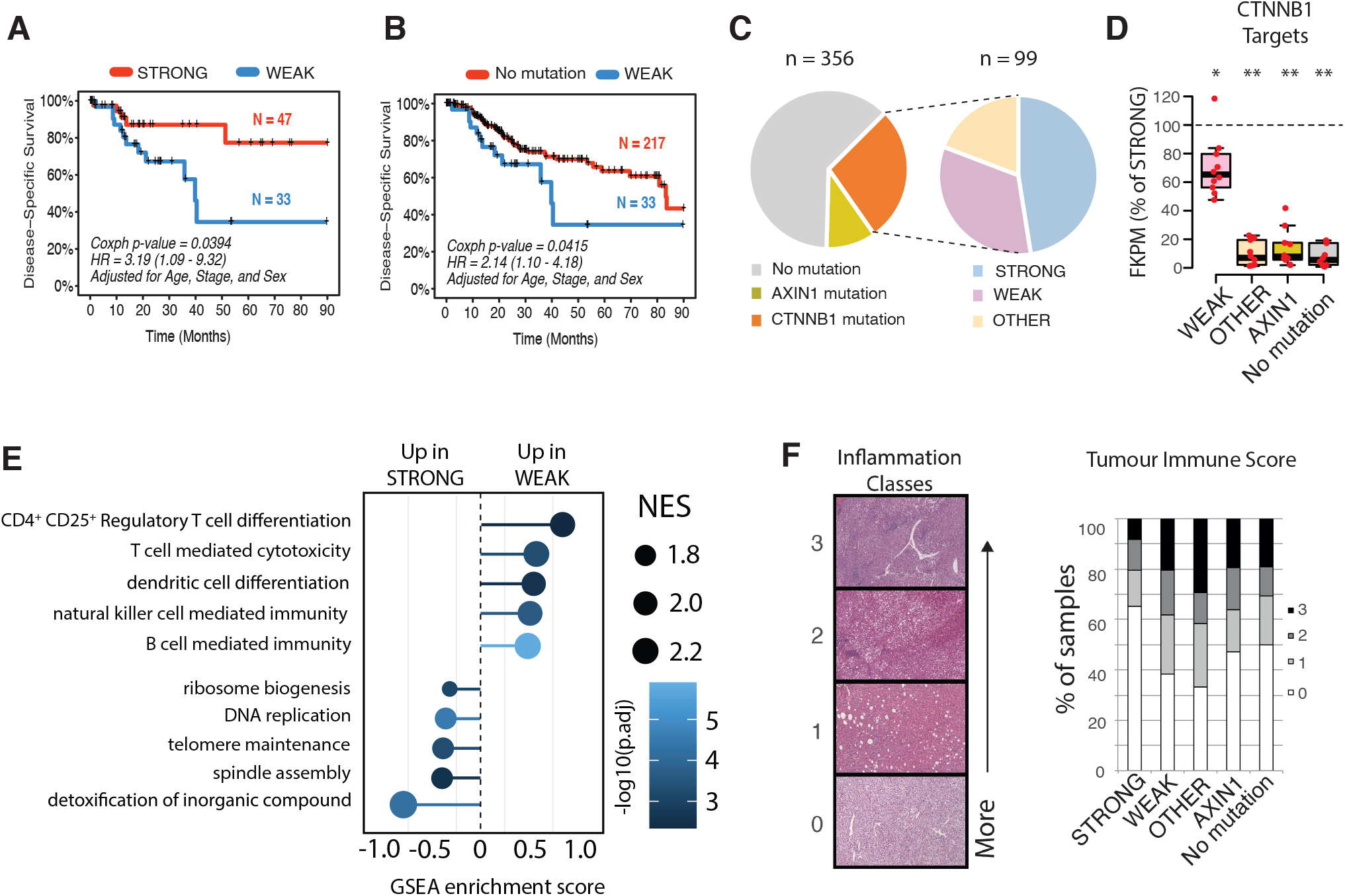
Mutation Effect Scores predict clinical and molecular features of hepatocellular carcinoma. **A & B** Differences in disease-specific survival between HCC patients from TCGA with (C) exon 3 mutations classed as STRONG versus WEAK (C) or exon 3 mutations classed as WEAK versus no mutation in *CTNNB1* or *AXIN1*. **C**. The proportion of HCC samples from the TCGA cohort with *CTNNB1* missense or *AXIN1* coding mutations (left). The *CTNNB1* missense mutations are then further divided into three categories (right). 18 further samples fell into >1 category so were excluded. TCGA Sample IDs are listed in Table S9 **D.** Expression of ß-Catenin target genes in HCC stratified by ß-Catenin pathway mutation status. Each red point represents the median expression value for one gene across all samples in the relevant patient group, expressed as a percentage of the median value for the same gene in the STRONG group (dashed horizontal line). Horizontal lines show the median value across all genes, boxes show the upper and lower quartiles and whiskers show the range.. * p < 4 e-03, ** p < 1 e-4 from negative binomial GLM. Gene level data are shown in Figure S6A. **E.** Enrichment of Gene Ontology terms in transcripts ranked among the most upregulated in pairwise comparisons between STRONG and WEAK *CTNNB1* mutant HCC. NES shows enrichment scores normalised to the size of the Gene Set. **F**. H&E tumour sections from TCGA were scored using an ordinal scale according to the level of inflammatory infiltrate from 0 (no visible immune cells) to 3 (diffuse or nodular aggregates of immune cells), then scores were compared across patients based on ß-Catenin pathway mutation status.

Although our mutational scanning assay measured the phenotype of 81% of *CTNNB1* missense mutations in the LIHC cohort, it did not assess all mutations implicated in the β-Catenin signalling pathway (Figure 5C). These include other missense mutations in *CTNNB1* that occurred outside the hotspot region (OTHER, n = 19), primarily at positions 335, 383 and 387 within armadillo repeats 5 and 6, which have been shown to perturb interactions with the destruction complex subunit APC (Liu et al. 2020). Moreover, a further 10% of the LIHC cohort had coding mutations in AXIN1 (n = 36): another destruction complex subunit and known HCC driver (Figure 4F, (Feng et al. 2012; Cancer Genome Atlas Research Network 2017)). We therefore explored the impact of these variants, which were ‘unseen’ by our mutational scanning assay, relative to WEAK and STRONG hotspot mutations on the CTNNB1 pathway in hepatocellular carcinoma. Based on published literature, we identified a set of 10 known ß-Catenin targets from among the top 50 upregulated genes in the transcriptomes of HCC tumours with any hotspot missense mutation, versus the NO MUTATION group. As expected, tumours with either WEAK or STRONG activating mutations in *CTNNB1* exon 3 expressed *CTNNB1* target genes at substantially higher levels compared to the NO MUTATION group (Figure S7A). Tumours with OTHER mutations in *CTNNB1*, or mutations in *AXIN1*, expressed these target genes at levels comparable to the NO MUTATION group (Figure 5D, Figure S7A). The OTHER group showed poor survival, similar to tumours with WEAK exon 3 mutations, whereas *AXIN1* mutations correlated with a more favourable outcome comparable to the STRONG group (Figure S6C). Thus, although mutations in *AXIN1* and *CTNNB1* armadillo repeats clearly contribute to HCC development, their effects on β-Catenin target gene expression are distinct from both WEAK and STRONG activating mutations within *CTNNB1* exon 3 ((Feng et al. 2012), (Abitbol et al. 2018)).

The differential expression of ß-Catenin targets in WEAK versus STRONG tumours was relatively subtle in comparison to the three other groups. Nonetheless, these genes showed significantly reduced expression in tumours from the WEAK versus the STRONG group (Mean 29% reduction, p < 0.001 (negative binomial GLM) Figure 4D, Figure S7A). The WEAK group therefore still robustly activates *CTNNB1* target genes, but to a degree that is intermediate between the STRONG activating group and tumours with no *CTNNB1* pathway mutation.

Transcriptome-wide, only modest differences were observed between HCC tumours with WEAK versus STRONG hotspot mutations (Figure S8, Table S4), despite the significant difference in survival between these groups (Figure 5A). However, gene set enrichment analysis identified cell proliferation-related terms as enriched among genes upregulated in the STRONG group (Figure 5E, Table S5); notably, the telomerase subunit and HCC driver *TERT* was among the most significantly upregulated genes (70% upregulated, adjusted p = 6.05 x 10^-7^, Table S4). *TERT* may be a direct transcriptional target of ß-Catenin regulation (Hoffmeyer et al. 2012), and activating mutations in the TERT promoter are among the most frequent somatic driver mutations in HCC (Cancer Genome Atlas Research Network 2017; Nault et al. 2014).

Numerous terms associated with lymphocyte infiltration were enriched among genes ranking among the most upregulated in tumours with WEAK versus STRONG exon 3 mutations (Figure 4H). It is now well established that β-Catenin signalling is a major pathway for immune escape in several tumour types, including HCC (Spranger et al. 2015; Ruiz de Galarreta et al. 2019), with potential implications for patient stratification and targeted therapy (Harding et al. 2019; Felden et al. 2021; Ruiz de Galarreta et al. 2019). Accordingly, canonical T cell transcripts were upregulated in tumours with WEAK versus STRONG activating *CTNNB1* mutations (Figure S7B), and a larger fraction of WEAK versus STRONG mutant tumours showed evidence of immune cell infiltration when histologically assessed using an ordinal scale (65 vs 38% with a score of 1 or greater, Fisher’s exact p = 0.0245, Figure 4I).

Overall, these data demonstrate that phenotypic scores measured in a cellular assay (Figure 2) provide novel insights into ß-Catenin regulation (Figure 3) and levels of oncogenic signal activation across tissues (Figure 4). They also correlate with important molecular and clinical features of hepatocellular carcinoma (Figure 5), which can be rationalised based on known pathways of ß-Catenin function (Ruiz de Galarreta et al. 2019; Hoffmeyer et al. 2012). Importantly, our novel mutational scanning approach using a cell-autonomous reporter assay readout (Figure 2, Figure S2) could predict non-cell autonomous phenotypes associated with the tumour microenvironment (Figure 5 & 4I, Figure S7B).

These insights were enabled by an experimental design with two advantages over previous studies that have addressed the phenotype of *CTNNB1* mutations. First, variants were introduced by genome editing at the endogenous chromosomal locus to avoid artefacts arising from ectopic expression. Second, the function of each variant was evaluated in parallel in the same population of cells, providing sensitivity to distinguish subtle differences in phenotype. Although our mutational scanning assay spanned just 2% of ß-Catenin amino acids, this covers over 75% of *CTNNB1*-mutant cancers in the COSMIC database (Table S1, (Tate et al. 2019)), including 74 different recurrent missense substitutions observed collectively in over 6000 tumours.

Using only experimentally determined mutational effect scores, we identified a subgroup of HCC patients characterised by relatively weak CTNNB1 pathway activation, immune infiltration and poor survival. Tumours with stronger activating mutations upregulated the *TERT* driver gene and tended to lack immune infiltration. The WEAK *CTNNB1* mutant subgroup is notable for the combination of immune infiltration and poor survival, because immune infiltration correlated with improved survival across the whole TCGA cohort (Figure S6D). However we note the similarities between the WEAK group and the ß-Catenin “non-excluded” immune subtype recently described by Montironi *et al* (Montironi et al. 2023).

Weakly activating exon 3 mutations, which our work has systematically defined for the first time, could provide both a novel mechanism and biomarker for this patient group and, in the future, help to guide strategies for personalised combination-based therapies. ß-Catenin activation is a known mechanism of immune escape in other tumour types, notably melanoma (Spranger et al. 2015), where exon 3 mutations also span a broad range of effect sizes (Figure 4D). Determining whether an optimal level of ß-Catenin signalling enables other tumour types to balance oncogene activation and immune evasion is now an important priority for future studies.

## Materials and Methods

### CRISPR design and cloning

All gRNAs were designed using the optimized CRISPR design webtool (http://crispr.mit.edu/), Benchling (https://benchling.com/) and Wellcome Sanger Institute Genome Editing (WGE) (http://www.sanger.ac.uk/htgt/wge/). The gRNAs were cloned into either pSpCas9(BB)-2A-GFP (Addgene Plasmid #48138) or pSpCas9(BB)-2A-mCherry as described previously (Ran et al. 2013) and sequence verified by Sanger sequencing using U6/F primer (5’-GAGGGCCTATTTCCCATGATTCC-3’).

### Cloning the Puro TK targeting vector

A backbone vector (WT β-catenin vector) was generated by amplifying 5.4 kb region of *Ctnnb1* intron1-intron6 using primers *5’*GGTTGATACTACCTTGAGTACTC*3’* and *5’*GATTCACAGGGCTGCTAGTG*3’*. The amplicon was cloned into PCR4 TOPO vector using TOPO TA cloning Kit (Invitrogen). Then, a Gibson cloning reaction was set up including WT β-catenin vector (amplified using primers *5’-*GTGAGGCTTTCTTTGTTGGC-*3’* and *5’*-GTCAAAAGGCAGAATGAAAACAG -*3’)*, Pu TKamplicon (*5’-*CTGTTTTCATTCTGCCTTTTGACCATAGAGCCCACCGCATCC-*3’* and *5’*-GCCAACAAAGAAAGCCTCACTACC GGGTAGGGGAGGCG -*3)* and Gibson Assembly master mix (NEB) following the manufacturer’s guidelines.

### Derivation and maintenance of TCF/Lef:H2B-GFP embryonic stem cells

Mouse ES cells were derived from the TCF/Lef:H2B-GFP mouse line (Ferrer-Vaquer et al. 2010) as described previously (Czechanski A. *et al,* 2014). The cells were then co-transfected with the Puro TK targeting vector and *Ctnnb1* sgRNA1-4 (Supplementary table 1) using lipofectamine 2000 (Invitrogen) following the manufacturer’s protocol. After Puromycin (1µg/ml) selection, individual clones were picked, grown and validated as heterozygously targeted by Sanger sequencing. Cells were then routinely maintained on gelatin coated flasks in GMEM (Gibco) supplemented with 10% FBS (GE Healthcare -HyClone), 2mM L-glutamine (Gibco), 1mM Sodium pyruvate (Gibco), 0.1mM MEM non-essential amino acids, 0.1mM β-mercaptoethanol (Gibco), 3µM CHIR99021 (Axon Medchem) and 1 µM PD0325901 (Axon Medchem) and Leukemia Inhibitory Factor (hereafter referred to as R2i media).

### Generation of the homology-directed repair template library

A double stranded DNA library was synthesized by Twist Biosciences, with each 200bp fragment encoding a single distinct amino acid (n = 20 per site) spanning codons 31 to 48 of Ctnnb1. Each fragment was flanked by *BbsI* recognition sites for cloning, and synonymous mutations enabling specific amplification of HDR edited alleles. Each fragment was then cloned individually into a β-catenin destination vector containing 5.5kb β-catenin intron1-6 sequence (5’-GGTTGATACTACCTTGAGTACTC-3’ and 5’-GATTCACAGGGCTGCTAGTG-3’) with two *BbsI* sites flanking the target region. After ligation, the reactions were transformed into Stbl3 competent cells and incubated overnight in a shaker at 37C. An equal volume of inoculum from each pool was then combined and used as starter culture for a single maxiprep plasmid isolation of the pooled HDR template library (Qiagen Maxiprep kit).

### Transfection of homology-directed repair template library and flow sorting

TCF/Lef:H2B-GFP reporter cells with heterozygous Puro TK knockin at the endogenous *Ctnnb1* locus were maintained in R2i media. A total of 200x10^6^ cells were transfected with the pooled HDR template library and Pu TK sgRNA 1-4 in twenty-six 6-well plates, using Lipofectamine 2000 (Invitrogen) following the manufacturer’s protocol. Small molecule enhancer L755507 was used at a concentration of 5µM. On day 3, R2i media was replaced with GMEM (Gibco) supplemented with 10% FBS (GE Healthcare - HyClone), 2mM L-glutamine (Gibco), 1mM Sodium pyruvate (Gibco), 0.1mM MEM non-essential amino acids, 0.1mM β-mercaptoethanol (Gibco) and Leukemia Inhibitory Factor, including FIAU at 0.2 µM to selectively kill cells that had not deleted the TK cassette. On day 5, a single cell suspension was generated using trypsin and the cells were sorted based on GFP intensity into 6 equally logged bins using BD FACS Aria III (BD Biosciences). In a parallel control condition, cells underwent an identical genome editing and selection procedure but were harvested without flow sorting (“pool” sample). The experiment was repeated twice. The number of cells sorted from each bin is listed in Table S7.

### DNA isolation and sample processing for deep sequencing

Genomic DNA was isolated using DNeasy Blood and Tissue kit (Qiagen) according to the manufacturer’s protocol. A first round of PCR was performed using a forward primer which annealed upstream of the homology arm (*5’*-GTGGACATCAGAGGACAACTTG-*3’*) and a reverse primer that annealed to a region containing the HDR edited allele (*5’*-TGTCAACATCTTCTTCTTCGGGA-*3’*). Entire DNA sample was amplified in several 30 cycle reactions using Q5 Hot Start High-Fidelity 2X Master Mix (NEB), the amplicons were digested with DpnI (thermo scientific), gel eluted, and pooled.

For library preparation for Illumina sequencing platform, a second round of PCR was performed to incorporate Illumina specific barcode, primer pad, linker and adapters. The reactions were performed in triplicate for 14 cycles using Q5 Hot Start High-Fidelity 2X Master Mix (NEB), pooled and purified using AMPure XP beads (Beckman Coulter).

### Retrieval of COSMIC data for all CTNNB1 mutant cancers (V94_38)

Targeted and genome-wide mutations were downloaded from https://cancer.sanger.ac.uk/cosmic/download, filtered by gene CTNNB1, from COSMIC release v94 (28 May 2021). Mutations labelled ‘Substitution - Missense’ were extracted, excluding one MNP p.I35_H36delinsSN.

### Read processing and calculation of mutational effect scores

Adapters were trimmed and paired ends merged with NGmerge (https://github.com/harvardinformatics/NGmerge) to produce single end reads, which were aligned with bwa mem v0.7.17 (Vasimuddin et al. 2019) to the 162bp CTNNB1 reference sequence. A set of reads with a single missense on-target mutation, and no other mutations, was generated. Reads that did not fully cover the region targeted for mutagenesis (58-117bp) were excluded, as were alignments with any of: indels anywhere, no mutations in the target region, mutations only outside the target region, multiple mutations in the target region, synonymous mutations only, or mutations resulting in a codon that was not in the repair template library. Remaining reads had precisely one missense mutation specified from the HDR template library.

Read counts were normalised within each of the six experimental bins and two control conditions by dividing the number of reads for each mutation by the total number of filtered reads in that bin, such that the sum of normalised counts for all mutations in the bin was equal to 1. To calculate enrichment values relative to the starting population, the normalised count for each mutation in each bin was then divided by the normalised count for the same mutation in cells that had undergone the same genome editing and selection procedure but had not undergone flow sorting based on GFP. Lastly, each of the 6 enrichment values for each mutation was multiplied by the mean GFP fluorescence value for all cells in the corresponding bin, and then the 6 values were added together to yield the mutational effect score (MES).

Replicates had high Pearson correlation (0.577-0.988) and were merged. Plasmid and unselected pool codon frequencies by position had Pearson correlation 0.626, and linear regression intercept 0.0002 and coefficient 0.9285.

### Variant Effect Predictor Comparisons

Variant effect predictor scores were obtained for all *CTNNB1* single amino acid substitutions spanning amino acids 31 - 48 from 50 different methods, using the same pipeline as described recently (Livesey and Marsh 2023). The Spearman correlations were then calculated between the outputs of each predictor and the MES values. Note that some predictors only output scores for missense variants possible by single nucleotide changes meaning that the correlations were calculated from fewer mutations; however, this has been shown to have little effect on overall correlations or relative predictor rankings (Livesey and Marsh 2020).

### AAIndex

To investigate amino acid properties potentially related to the effects of mutations at I35, we downloaded all 566 indices from the AAIndex database (Kawashima et al, 2008). The Spearmann correlation was calculated between the values for each amino acid and the MES for each of the 19 substitutions at I35. These correlations are provided in Table S8. Peptide sequences for previously reported βTrCP-dependent degrons were extracted from Table S1 of (Low et al 2014), then filtered for a precise match to the DSGX motif spanning positions 32 – 35 of CTNNB1 (n = 28 peptides, detailed in Table S4).

### Stratification and survival analysis of the TCGA Hepatocellular Carcinoma Cohort

Samples from the TCGA LIHC cohort (n = 370) were filtered into one of six groups (Table S9): 1) Those with a hotspot missense mutation classified as STRONG (MES > 18000, n = 47); 2) Those with a hotspot missense mutation classed as WEAK (MES < 18000, n = 33); 3) Those with a *CTNNB1* missense mutation outside the hotspot region (n = 19); 4) Those with any coding mutation in *AXIN1* (n = 36); 5) Those without any coding mutation in *CTNNB1* or *AXIN1* (n = 217); 6) Those with deletions and/or complex mutations in *CTNNB1*, those falling into more than one of groups 1 to 4, and those lacking survival and/or RNA-Seq data (n = 18). Group 6 was excluded from further analysis. For stratification of exon 3 missense mutations into WEAK and STRONG groups based on EVE predictions, a threshold of 0.75 was selected to yield groups of almost identical size relative to the MES-based stratification.

The impact of CTNNB1 mutation strength on disease-specific survival (time between diagnosis and death as a result of HCC) was assessed using the R Survival library.

Multivariable Cox proportional hazards regression models were fitted, adjusting for the patient’s age and stage at diagnosis, as well as their sex. Similarly, Cox proportional hazards regression models were also fitted comparing CTNNB1 EVE scores (STRONG versus WEAK), histological immune scores (0 versus 1, 2, or 3), and AXIN1 coding and CTNNB1 missense mutation subdivisions (STRONG, WEAK, OTHER). Follow-up information, including disease-specific survival, was available for 362 of the HCC TCGA patients, of which 78 were deceased by the time of last follow-up (right-censored to 90 months). All results were reported as hazard ratios (HR) with corresponding 95% confidence intervals (95% CI).

### Gene Ontology

Gene Set Enrichment Analysis (GSEA): GSEA was performed to investigate the functional enrichment of the differentially expressed genes (DEGs) identified from our RNA-sequencing data. The ranked list of DEGs was generated based on the DESeq2 statistics, which takes into account both fold-change and p-value information. GSEA was performed on the Gene Ontology (GO) database, which consists of three structured, controlled vocabularies. The enrichment analysis was carried out using the ClusterProfiler (4.6.2) package in R (version 4.2.0) (Yu et al. 2012b), using the “gseGO” function with default parameters.

### Histological Scoring of Immune cell infiltration

Whole-slide images of haematoxylin and eosin-stained formalin-fixed, paraffin embedded sections (‘Diagnostic Slide’) of the TCGA LIHC cohort were viewed using the NCI GDC Data Portal slide viewer; cases where only a frozen section image (‘Tissue Slide’) was available were not evaluated. Each case was scored by an expert consultant liver histopathologist and National Liver Pathology External Quality Assurance scheme member working at the national liver transplant centre (TK) blinded to all new experimental data and without accessing any additional TCGA data. Intratumoral inflammation was scored using H&E morphology alone using a 4-point ordinal scale (Joung et al. 2018): 0, no inflammation; 1, scattered inflammatory cells; 2, focal aggregates of inflammatory cells; 3, diffuse or nodular aggregates of inflammatory cells.

### Differential expression analysis

STAR transcript quantification for TCGA-LIHC samples (n=361) was downloaded from the GDC Data Portal. Differential expression of samples in each category compared to weak MES missense mutation was calculated with DeSeq2 v1.34 (Love et al. 2014) using shrunken log-fold changes. Gene Set Enrichment Analysis (GSEA) was performed to investigate the functional enrichment of the differentially expressed genes (DEGs) identified from our RNA-sequencing data. The ranked list of DEGs was generated based on the DESeq2 statistics, which takes into account both fold-change and p-value information. GSEA was performed on the Gene Ontology (GO) database, which consists of three structured, controlled vocabularies. The enrichment analysis was carried out using the ClusterProfiler (4.6.2) package in R (version 4.2.0), using the “ gseGO” function with default parameters (Yu et al. 2012a). For the analysis of CTNNB1 target genes, 10 known targets (based on published literature) were identified among the 50 most upregulated transcripts in STRONG versus NO MUTATION tumour groups from the TCGA LIHC dataset. We modelled expression as predicted by gene identity, LIHC subgroup and their interaction using negative binomial models (R package MASS). We then obtained p-values comparing expression in each LIHC subgroup to “strong” as the baseline using marginal means testing implemented in the R package emmeans.

### Deriving nucleotide-level TNC scores from LIHC and UCEC exome data

Exome sequence variants from TCGA-LIHC (n=375) and TCGA-UCEC (n=404) were downloaded from the GDC Data Portal (https://portal.gdc.cancer.gov/). Variants called by at least 2 of the 4 provided workflows (MuSE, MuTect2, SomaticSniper, VarScan2) were retained. Cases with one or more missense mutation in the 31-48aa target region were extracted (TCGA-LIHC n=82, TCGA-UCEC n=104), and the SNPs from these used to generate tri-nucleotide mutation frequencies with SomaticSignatures (Gehring et al. 2015). The results shown here are based upon data generated by the TCGA Research Network: https://www.cancer.gov/tcga.

### Model to convert trinucleotide scores to mutational likelihood scores

All possible mutational paths from one codon to another, one nucleotide change at a time, were generated for all codon pairs. The mutational likelihood score (MLS) of each path was calculated as ∏_𝑖_(𝑓𝑟𝑒𝑞_𝑖_) where 𝑓𝑟𝑒𝑞_𝑖_ is the tri-nucleotide context mutational frequency of the 𝑖^𝑡ℎ^ step of the path. The MLS of each codon change in every possible tri-nucleotide context is the sum of all paths between the two codons, similarly the MLS for each amino acid change in its tri-nucleotide context is the sum of its codon change scores. Worked examples are shown in Figure S5B and Table S10, and MLS scores calculated separately from TCGA-LIHC and UCEC are listed in Table S11. For statistical testing, we modelled observed mutation frequencies as the result of mutational likelihood score in the same tumour type, and using a negative binomial GLM, implemented in the R package MASS. We compared these fits to null models fitting only the mean using likelihood ratio tests.

### Availability of code, data and biological materials

Custom scripts used for data analysis in this manuscript are available at https://git.ecdf.ed.ac.uk/igmmbioinformatics/betacatenin-saturation-screen. Tcf:Lef:H2B:GFP mouse embryonic stem cells are available from the authors upon request.

## Acknowledgements

PH acknowledges grant support from the MRC (MR/M010341/1) and BBSRC (BB/J004316/1). AW acknowledges grant support from the MRC (MC_PC_21040) and from an MRC Unit Award to the MRC Human Genetics Unit. We thank Greg Kudla and Ian Adams for advice and discussions.

## Supplementary Tables

**Table S1**

**Details of COSMIC tumours with *CTNNB1* mutations**

**Table S2**

**Mutational Effect Scores for all hotspot mutations.**

**Table S3**

**Correlation coefficients for Mutational Effect Scores and in silico Variant Effect Predictors**

**Table S4**

**High confidence ß-TRCP docking sites**

**Table S5**

**Differential expression analysis on RNA-Seq data from stratified HCC tumours**

**Table S6**

**Gene Set Enrichment analysis on differentially expressed genes**

**Table S7**

**Cell numbers collected from FACS following saturation genome editing**

**Table S8**

**Spearmans ρ values for MES measured against indices from AAindex**

**Table S9**

**Details of TCGA IDs for Stratified Hepatocellular Carcinomas**

**Table S10**

**Worked example calculation of a mutational likelihood score**

**Table S11**

**Mutational likelihood scores for all hotspot missense mutations**

---

**Figure S1:**
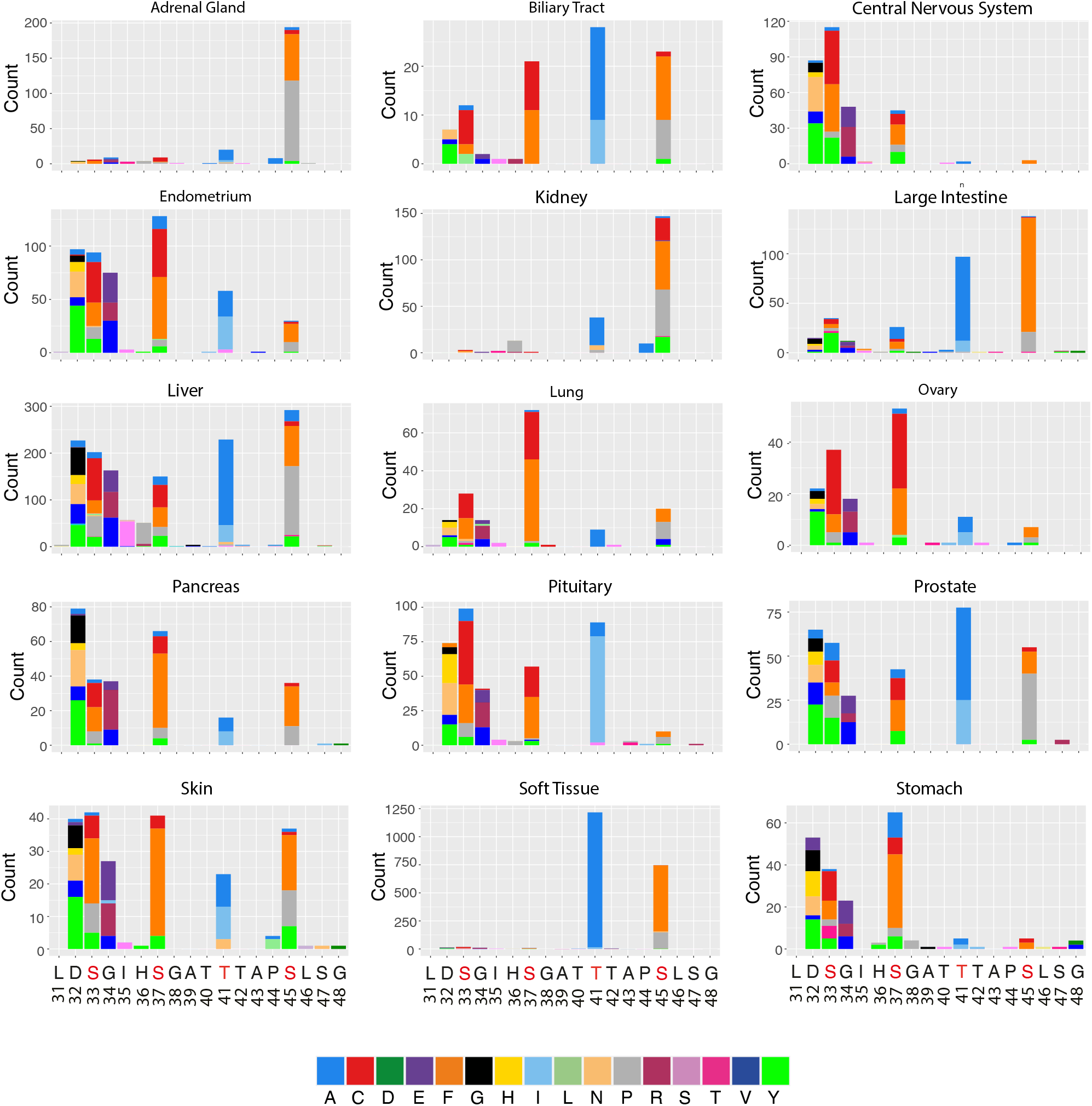
Tissue-specific mutation profiles at the *CTNNB1* degron in human cancer. Histograms show distinct distributions of mutations within the CTNNB1 mutation hotspot for COSMIC tumours filtered by different primary tissue sites. Only tumours with >100 mutations within the exon 3 hotspot are shown. Plots showing data for Adrenal gland, Central Nervous System and Skin are the same as those shown in Figure 1D.

**Figure S2:**
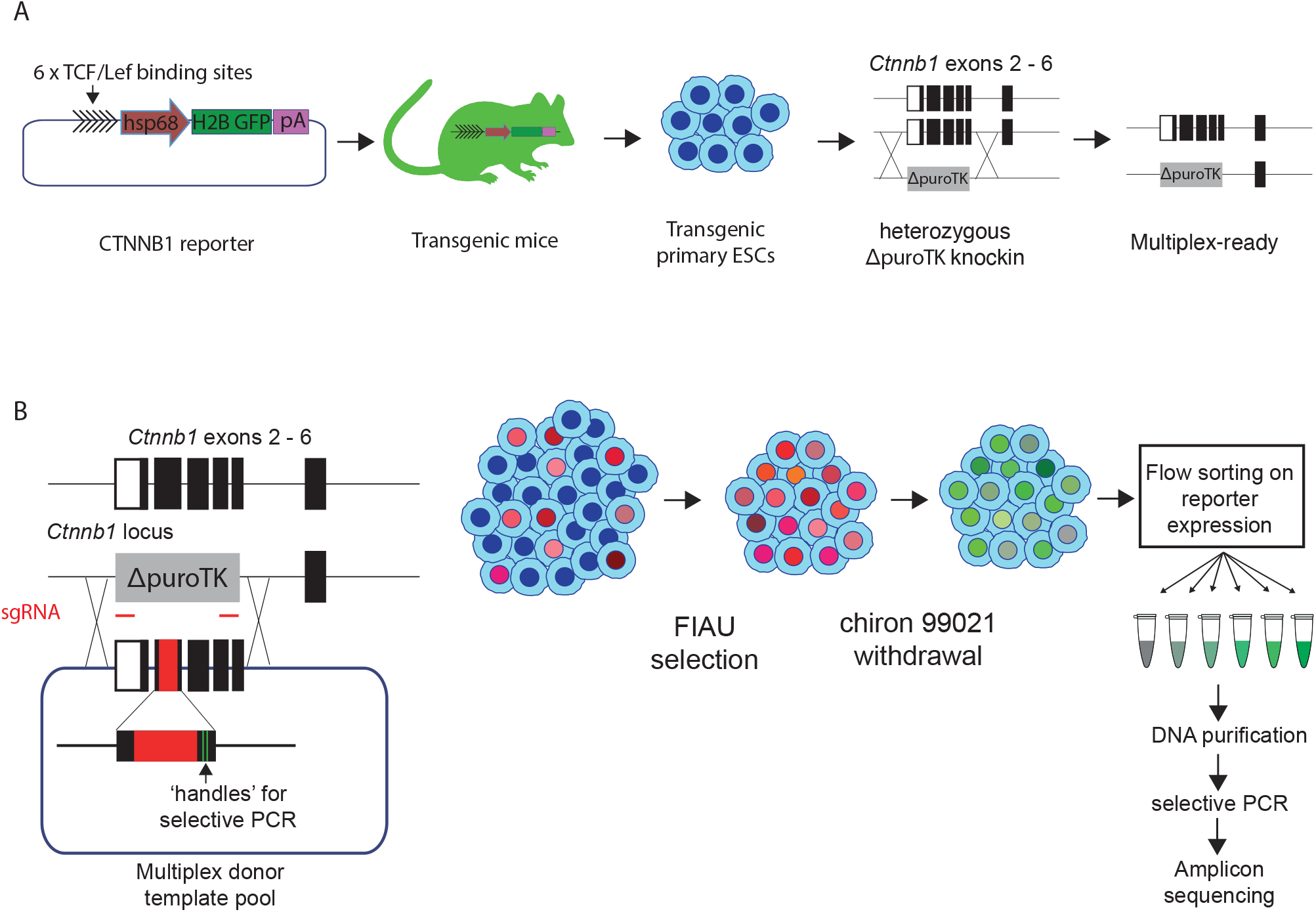
A mutational scanning assay to quantify ß-Catenin activation resulting from exon 3 mutations. **A.** The ß-Catenin transcriptional reporter construct used in this study, as previously described in (Ferrer-Vacquer 2010). Tandem copies of Tcf/Lef binding motifs were inserted upstream from a core hsp60 promoter driving expression of H2B:GFP, then this construct was integrated randomly in to the genome. Primary embryonic stem cells were derived from these mice, and a region spanning exons 3 – 6 of *Ctnnb1* was replaced with a ΔpuroTK selection cassette on one of two alleles to generate ‘multiplex ready’ cells **B.** A plasmid library was generated for homology directed repair, encoding each of 342 possible single amino acid substitutions spanning codons 31 to 48 of CTNNB1, together with silent mutations to allow selective PCR amplification of edited alleles from genomic DNA. This was transfected into multiplex-ready mESCs together with two sgRNAs that cut in an allele-specific manner, on either side of the selection cassette. Transfected cells were then cultured in the presence of FIAU to kill those in which the selection cassette had not been removed. Multiplex ready cells were routinely cultured under 2i conditions (Qing *et al* 2008), but The GSK3ß inhibitor was withdrawn 2 days before transfection to allow CTNNB1 signalling to return to a baseline state. Cells were then subject to fluorescence activated cell sorting based on the level of GFP reporter expression, before extraction of genomic DNA, amplification of the CTNNB1 exon 3 region by PCR, and Illumina sequencing. Further methodological information is detailed in the Methods section.

**Figure S3:**
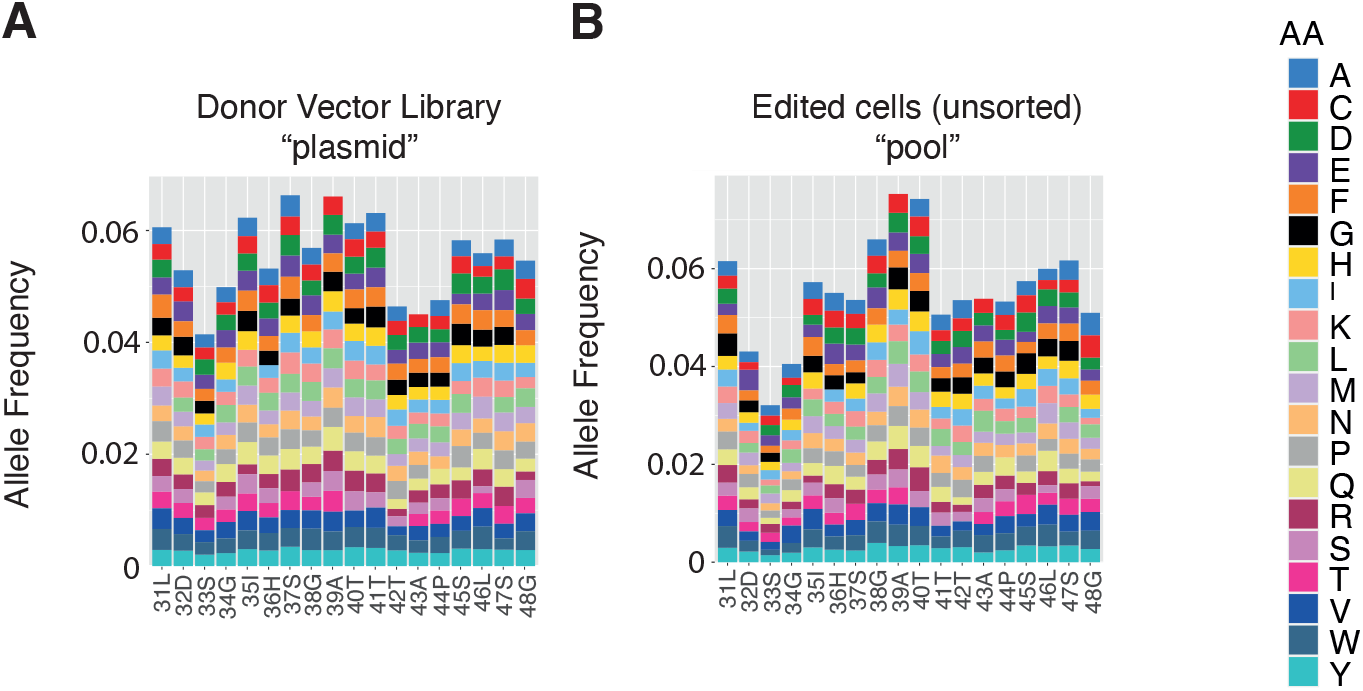
Allele frequencies in control sequencing libraries. The frequency of individual missense mutations in **A.** the “plasmid” library used for homology-directed repair, and **B.** in the cellular “pool” after editing and selection but before sorting based on GFP expression. The colour scheme used to represent different missense mutations is shown to the right. The “pool” sample was used to determine enrichment of individual mutations in each fluorescence bin during the calculation of mutational effect scores (Figure 2F).

**Figure S4:**
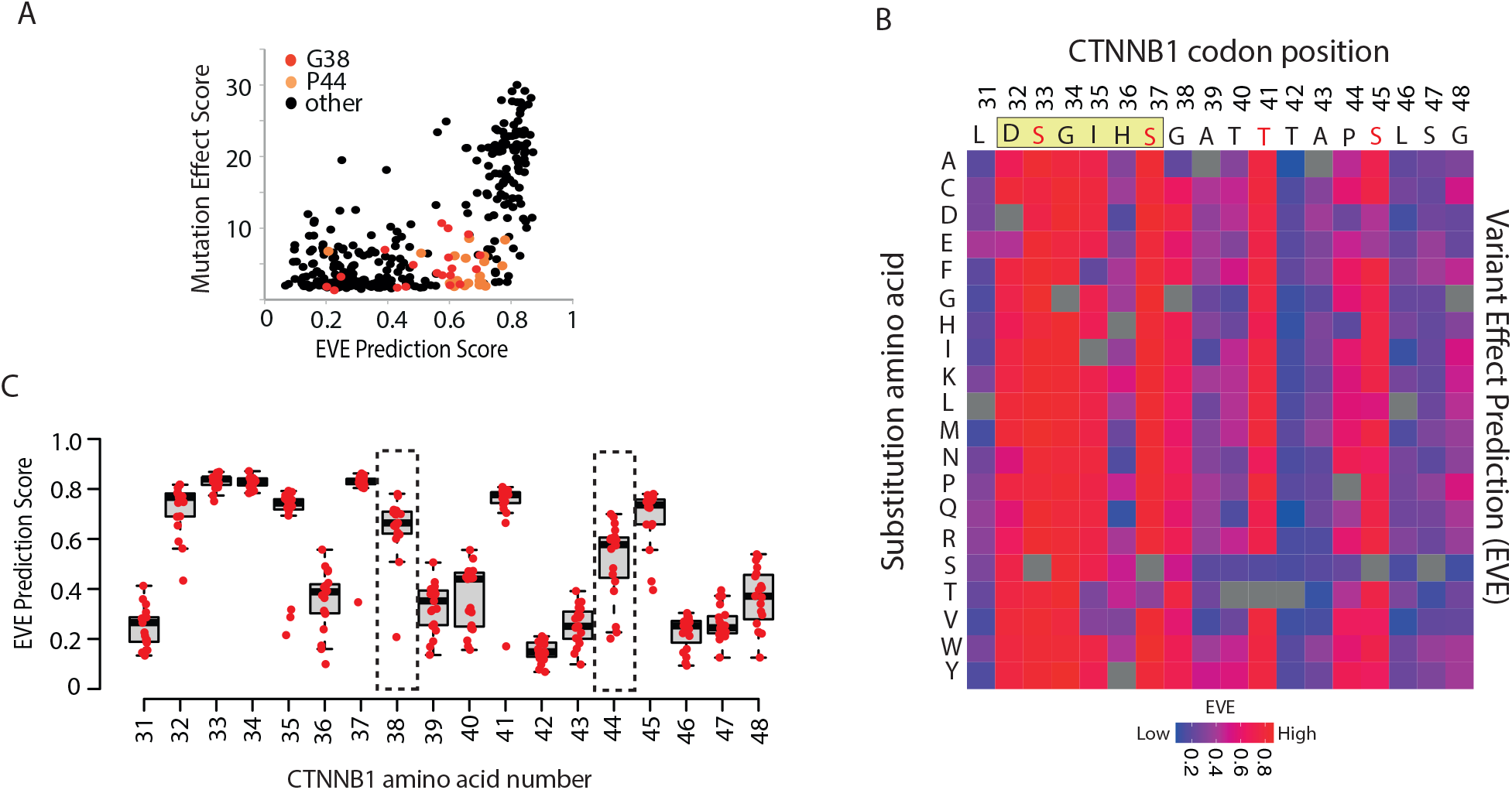
Comparison of Mutation Effect Scores with a top-performing *in silico* variant effect predictor. **A.** Dotplot to show correlation between mutational effect scores from this study, and *in silico* mutational effect predictions by the EVE tool (Frazer et al. 2021). Points corresponding to variants at positions G38 and P44 are highlighted. **B**. Heatmap representation of *in silico* Mutational Effect Predictions for all possible amino acid substitutions by the EVE tool. *CTNNB1* codon positions are indicated along the top, with the ß-TRCP docking motif highlighted yellow and phosphosites shown in red text. **C.** The distribution of EVE prediction scores for each codon position of *CTNNB1*. Horizontal lines show the median value, boxes show the upper and lower quartiles and whiskers show the range. Positions G38 and P44 where scores are discordant with cancer mutation frequencies (Figure 1) and MES values (see Figure 2F, Figure 3A) are highlighted.

**Figure S5:**
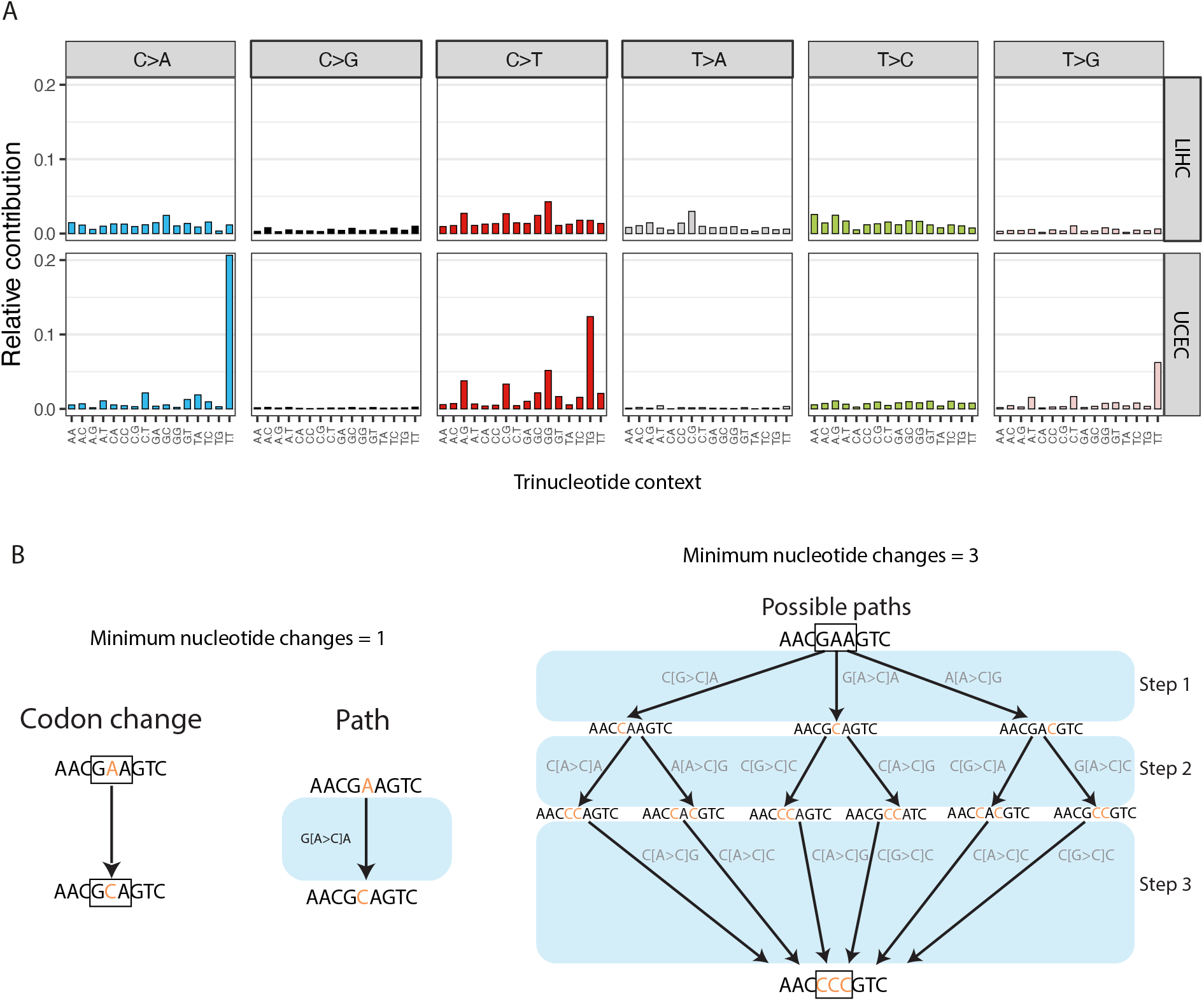
Calculating Mutational Likelihood Scores for amino acid substitutions in the exon 3 hotspot. **A.** Histograms show the relative frequency of all possible nucleotide substitutions, calculated in all possible trinucleotide contexts (n = 96), from exome sequencing data from TCGA Hepatocellular Carcinoma (LIHC – top, n = 82) and Uterine Endometrial Carcinoma (UCEC – bottom, n = 104). Only tumours with exon 3 *CTNNB1* hotspot mutations were used. **B.** Model to illustrate the process through which nucleotide-level mutation probabilities shown in panel A are converted to amino acid-level mutational likelihood scores (MLS). Amino acid substitutions can be reached by a minimum of either 1, 2 or 3 steps. The left panel shows a simple example where only a single nucleotide change is required to reach a destination codon. Where >1 change is required (example in the right panel), the probability of alternative paths is combined. In all cases, the probability of all paths to different triplets encoding the same amino acid are integrated to generate the MLS. Further details are provided in the Methods section.

**Figure S6:**
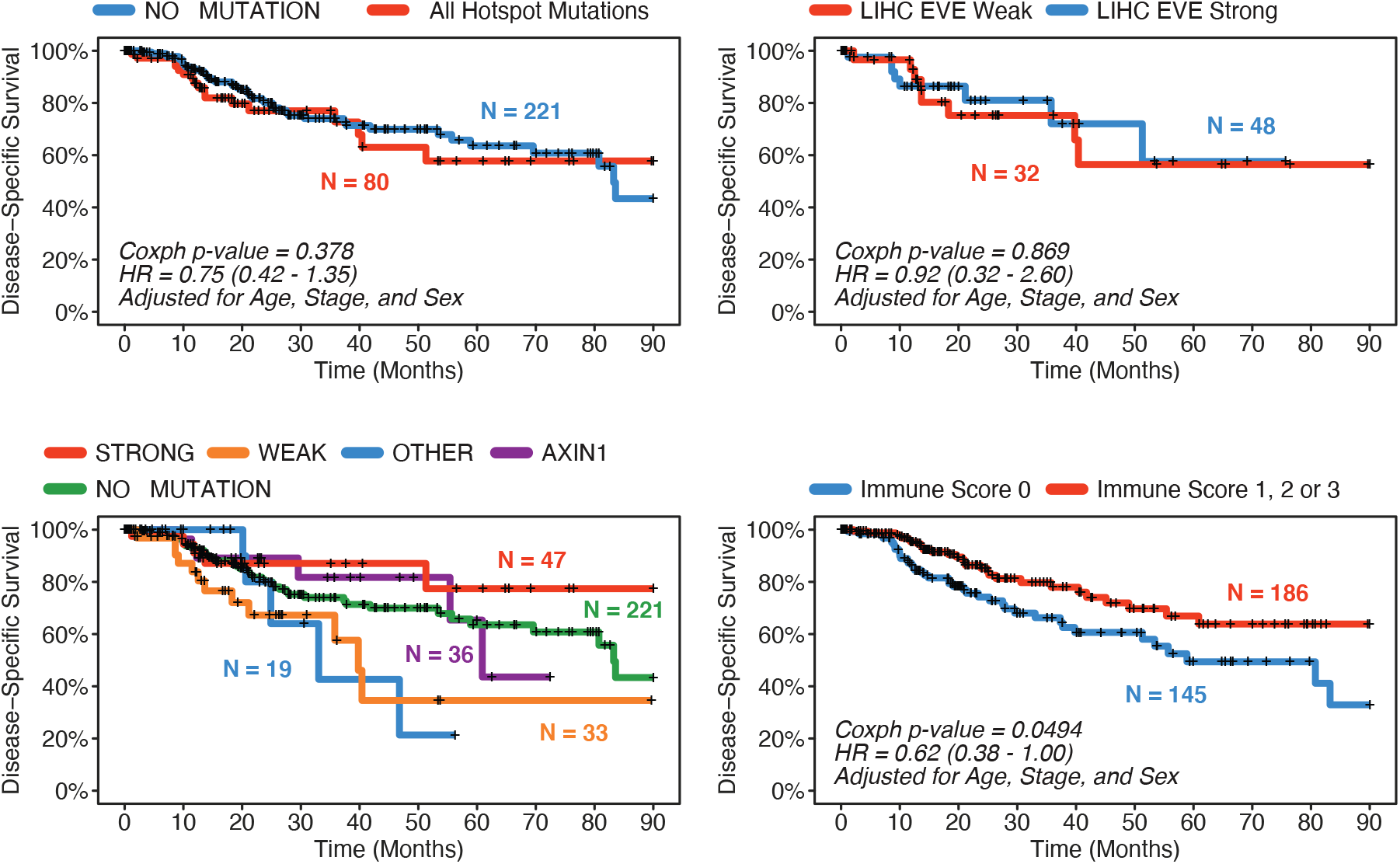
Survival analysis of Hepatocellular Carcinoma patients from TCGA. Kaplan Meier plots shows differences in disease-specific survival between HCC patients from TCGA with (**A**) any missense mutation within the *CTNNB1* exon 3 hotspot versus no mutation in *CTNNB1* or *AXIN1*; (**B**) exon 3 hotspot mutations classified into WEAK or STRONG categories based on variant effect predictions using the EVE tool (Frazer *et* al. Nature 2021); (**C**) All categories of HCC shown in Figure 4F; (**D**) histological immune scores of 1 or greater, indicating at least some immune infiltration evident on H&E-stained tissue sections, versus histological immune scores of zero.

**Figure S7:**
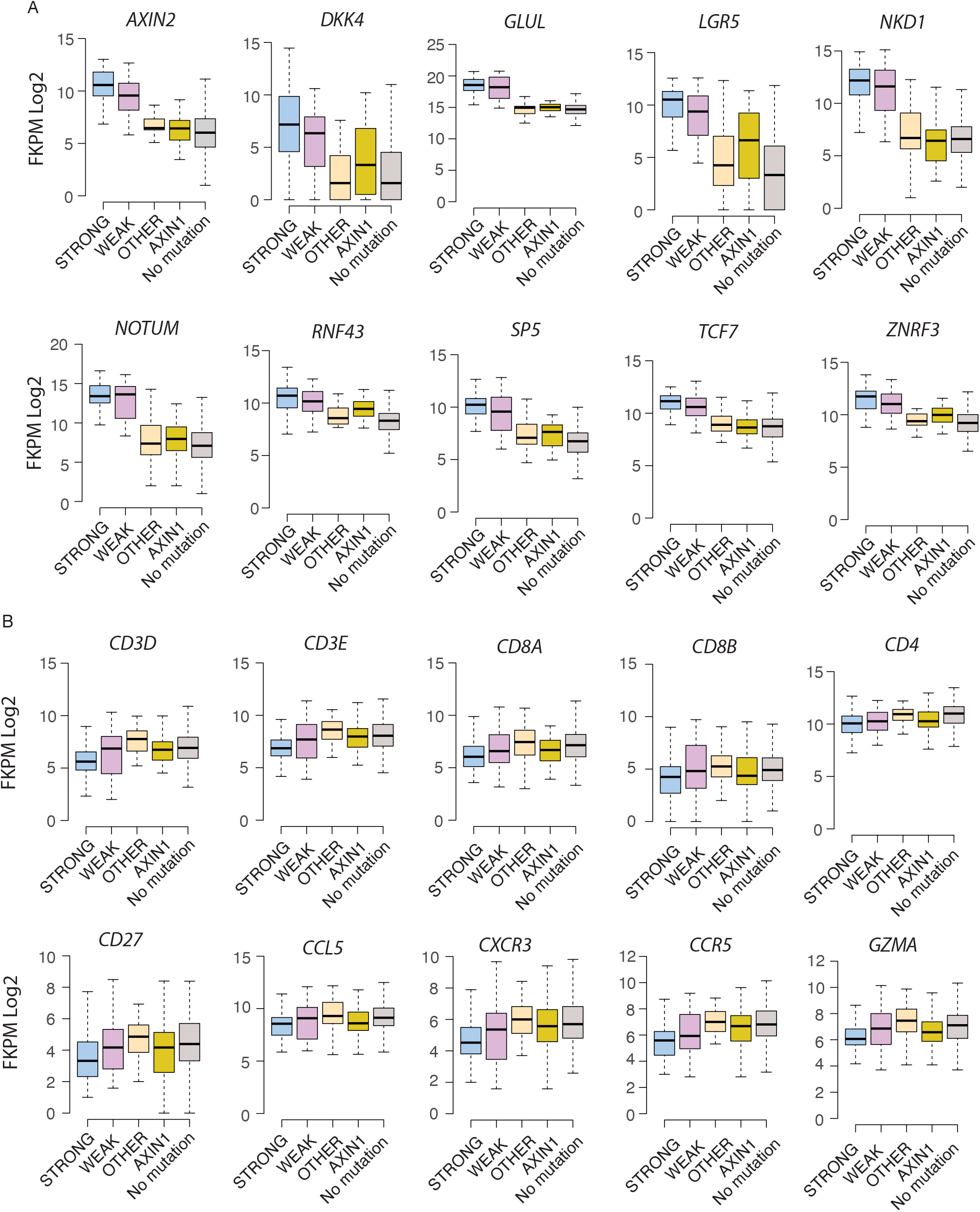
RNA-Seq expression analysis of individual genes in Hepatocellular Carcinoma. **A.** Expression of 10 canonical ß-Catenin target genes in TCGA Hepatocellular Carcinoma samples stratified as indicated in Figure 5C. Median expression values from each gene in each group are shown in Figure 5D. Horizontal lines show the median value, boxes show the upper and lower quartiles and whiskers show the range. **B.** Expression of 10 T cell signature genes in TCGA Hepatocellular Carcinoma samples stratified as indicated in Figure 5C.

**Figure S8:**
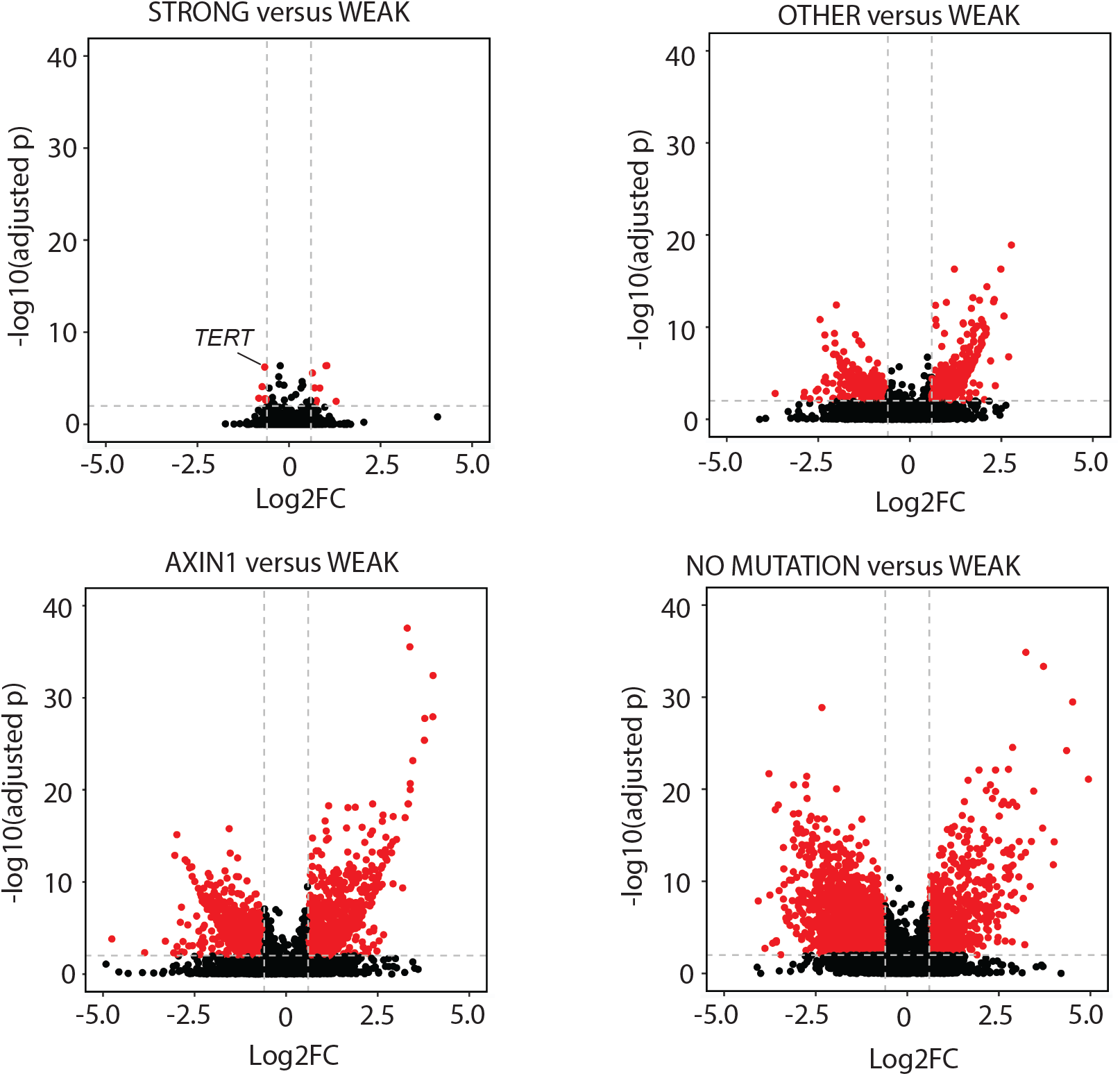
Transcriptomic differences between HCC samples grouped by ß-Catenin pathway mutation status. Volcano plots show the Log2 fold change and -Log10 adjusted p-values for differentially expressed genes in transcriptome comparisons between tumours with WEAK mutations in the *CTNNB1* hotspot region and other classes shown in Figure 5C. Dotted lines indicate significance thresholds (Log2FC greater than 0.6, adjusted p value < 0.01), and points coloured red denote genes which pass these thresholds. The values for individual genes are shown in Table S4, and Gene Ontology term enrichments in Table S5.

